# Fragment-Guided New Therapeutic Molecule Discovery and Mapping of Clinically Relevant Interactomes

**DOI:** 10.1101/2025.07.09.663848

**Authors:** Austė Kanapeckaitė, Sarper Okuyan, David James Wagg, Jan Koster, Ligita Jančorienė, Indrė Sakalauskaitė, Birutė Brasiūnienė, Andrea Townsend-Nicholson

## Abstract

Therapeutic interventions for complex diseases depend on the targeted modulation of key pathological pathways. While growing clinical needs continue to drive advancements in the drug discovery space, current strategies primarily rely on searching large volumes of chemical data without addressing the specific contributions of molecular features. Moreover, both clinicians and researchers recognize the need for improved drug discovery methods and characterization that could aid in clinical strategy selection. To address these challenges, we propose a new perspective on targeted therapy development as well as interactome mapping, utilizing molecular fragments. The present study focuses on therapeutic areas that represent emerging targets, namely JAK2 and GLP-1R, both of which have broad clinical potential. We developed a new self-adjusting neural network that enabled us to discover novel therapeutic candidates with improved in silico binding profiles, gain additional insights into drug-target binding that were not previously reported, and identify new metabolic trajectories. Importantly, our work revealed that even a small compound library can effectively generate lead candidates, expediting the search and exploration process. In addition, the fragment-guided bridging of chemical and biological spaces has revealed new opportunities for drug repurposing efforts and a means of improving the prediction of side effects. We concluded our study with insights into the recent high-profile clinical trial failure of danuglipron and how this could have been prevented with our methodology. Thus, building a robust in silico pipeline with integrated screening data can significantly reduce costs and guide therapy adoption. Furthermore, our proposed strategy highlights promising avenues for the discovery of new therapeutics and the development of clinical interventions.

**TOC Graphic:** 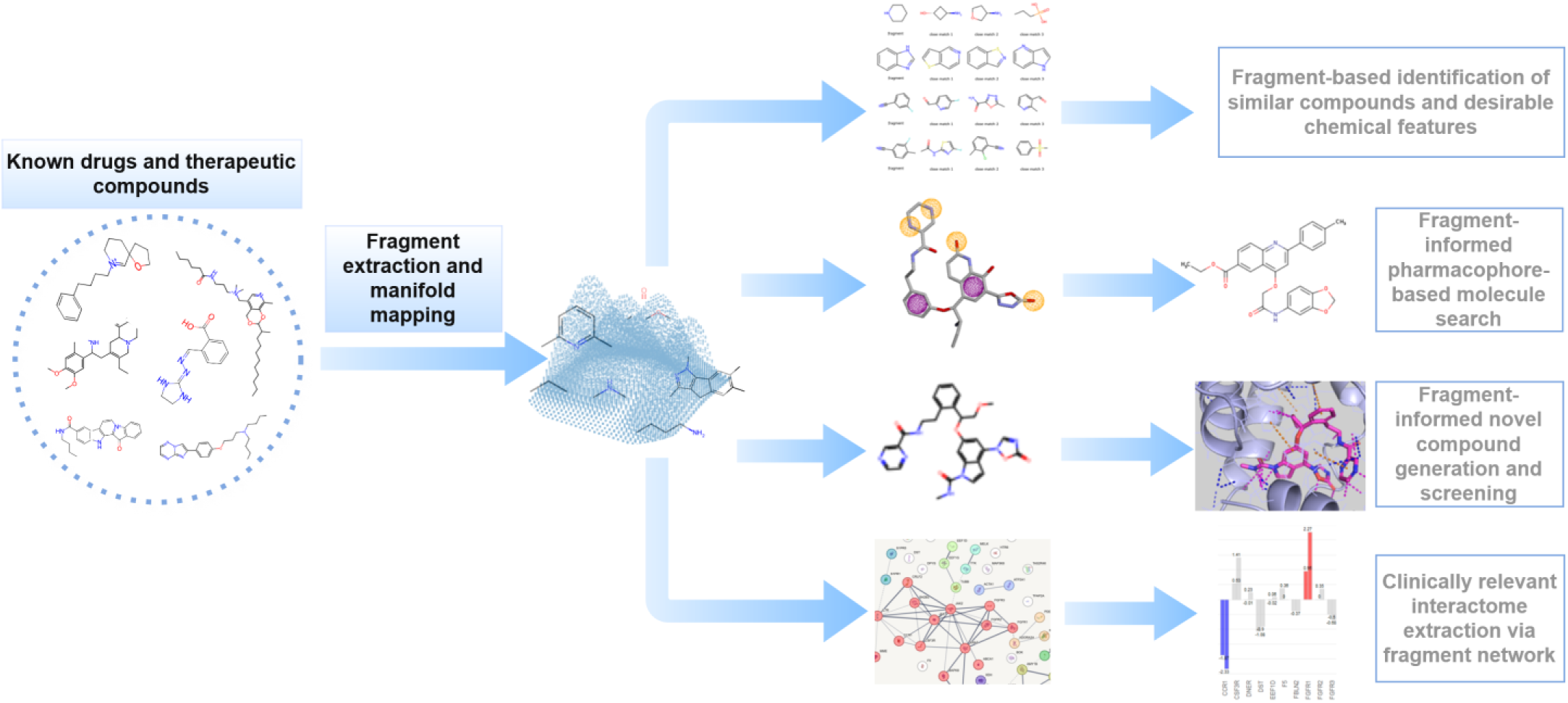

## Introduction

The drug discovery process can follow several trajectories, depending on the search strategy. Whether the choice is to start with a specific target or a compound library, the goal is to identify therapeutic candidates with promising activity. This involves a gradual refinement of hits to lead compounds that exhibit favorable drug-like properties (1, 2). Moreover, traditional high-throughput methods have been augmented with computer-aided drug design (CADD) to expedite screening and detection. Nevertheless, the chemical space is estimated to be in the order of 10^64^ , rendering the exploration computationally intractable when using traditional methods (2). Consequently, new computational strategies are being developed to address the combinatorial complexity of iterating through candidate molecules. Despite the wealth of tools, limitations remain, as a more targeted drug search is needed to sift through chemical variation and to detect features of therapeutically relevant, drug-like compounds (2, 3). To tackle these challenges, we need to leverage several observations that emerged in the clinical space. Firstly, drugs and active compounds often exhibit polypharmacological effects (i.e., modulation of multiple targets) (4). This suggests that a compound of interest might have several functional groups that either separately or in combination influence specific therapeutic outcomes across several targets. Secondly, drug-target stoichiometry and response variability indicate that both the drug’s efficacy and potency might be related to the functional group composition (5). Given these insights, we propose a targeted exploration guided by fragmenting active compounds and creating a functional probabilistic space mapping that can be explored and juxtaposed to other readouts (e.g., structure-activity relationships, target networks, disease interactome, and others). Using this method, instead of trying to group entire compounds — which could prevent us from capturing therapeutically meaningful subgroups — the fragmentation allows us to map the chemical interactome across different therapeutics (Fig.1). Specifically, we focus on active (or assumed to be active) subsets to create ‘probability valleys’ that can enable targeted feature search and the generation of novel drug candidates via the fragment assembly. Our work demonstrates that even a small library of around 10,000 molecules is sufficient to generate highly diverse compounds when compared to known drugs. This contrasts with the expanding library investments and scale (billions of molecules), which studies have shown can result in missing drug-like candidates (6). In addition, we overlaid fragments with known drug targets and captured additional therapeutic trajectories as well as flagged potential side effect contributors. Therefore, switching from brute-force approaches to rational mapping and design could not only help uncover previously overlooked chemical spaces but also improve cost-effectiveness and speed of the discovery process. To test these ideas, we focused on therapeutic areas with both unmet needs and broad clinical impact potential.

**Figure 1.**
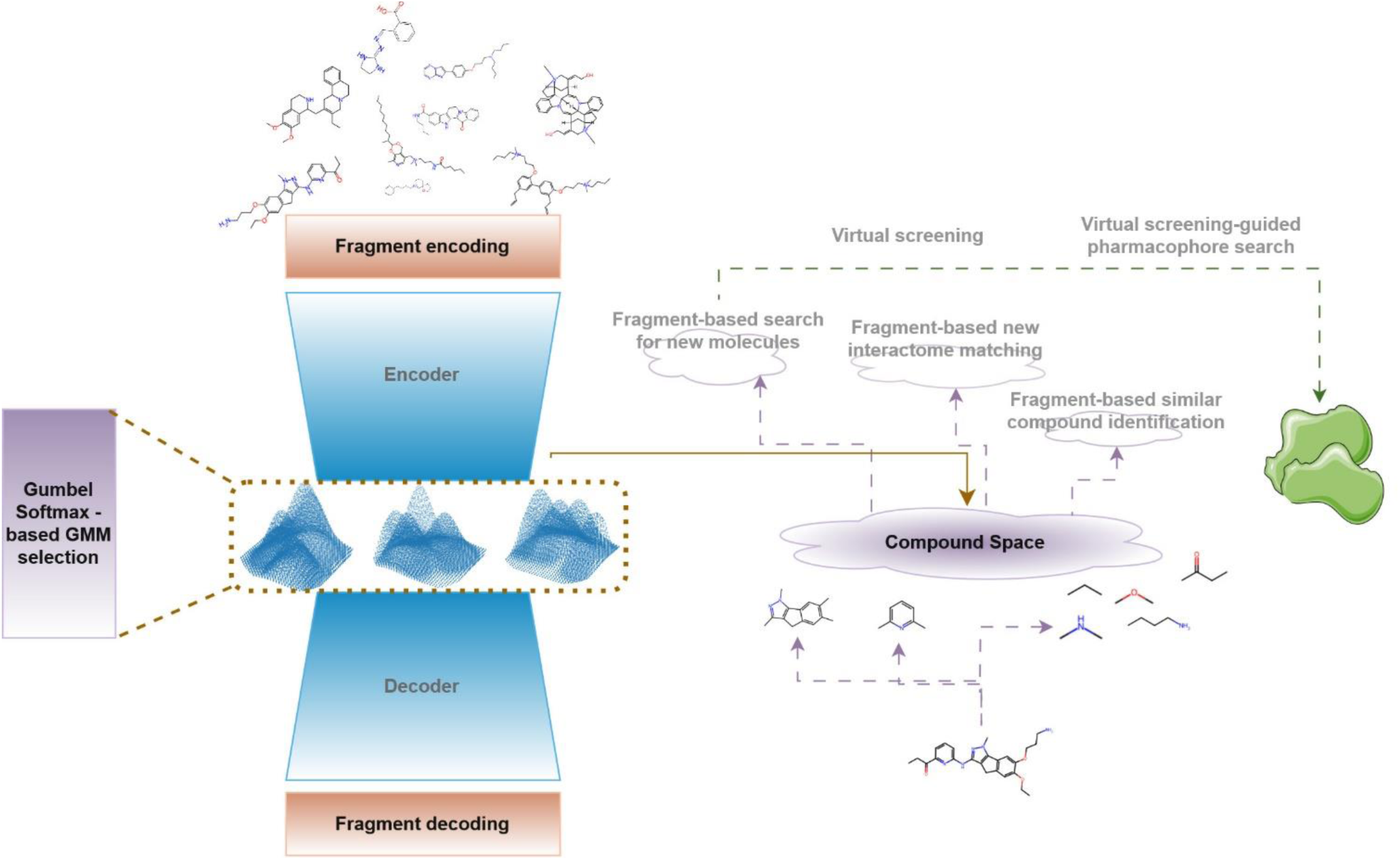
Model and workflow overview. The analysis begins with adaptive learning-based Gaussian mixture latent space mapping of compound fragments. The mapping is used to identify fragments for new molecule generation and relevant interactome identification.

For a successful mapping campaign, we integrated several cutting-edge methods, namely positional encoding (PE) with Laplacian eigenvectors (7, 8) and the graph isomorphism convolution layer (GIN) (9), into an adaptive Gaussian mixture (GM) variational network.

Permutation equivalence, uniqueness, and distance awareness with respect to the graph topology are some of the key features of Laplacian eigenvectors, providing a meaningful coordinate system and preserving both the local and global graph structure for compounds of interest. Thus, in our approach, Laplacian PE is used to update fragment features before passing them into a GIN layer for molecular feature embedding. The number of clusters in the variational space is adaptively learned with a Gumbel softmax layer (10). Consequently, the low-dimensional GM embedding of molecular fragments enables the learning of data point distributions and allows the network to self-adjust based on the inputs (11) (Fig. 1).

Because 35% of approved drugs target G protein-coupled receptors (GPCRs) and another major class represents kinase inhibitors, we selected therapeutic scenarios focusing on these protein families and their associated drugs (12, 13). The first candidate was Janus kinase (JAK), which, after phosphorylation, activates the signal transducer and activator of transcription (STAT), initiating a cascade of intracellular and extracellular signaling (14). To investigate JAK modulation, ruxolitinib (INCB018424) was chosen since it is a selective type I kinase inhibitor of JAK1 and JAK2. The drug’s action reduces cellular proliferation, lowers cytokine plasma levels, and induces apoptosis (15). Due to intricate regulatory networks, modulating JAK1/JAK2 signaling shows potential in treating not only myeloproliferative neoplasms (MPNs), but also inflammation, as well as glucose and lipid dysregulation (14, 15). Furthermore, recent studies have highlighted JAK/STAT involvement in obesity and type 2 diabetes mellitus. Thus, chronic inflammatory conditions could be responsive to JAK inhibitors (JAKi), offering broad applications in clinical care from immune regulation and oncology to surgery and tissue healing (16, 17, 18). The second target was the glucagon-like peptide-1 receptor (GLP-1R), paired with an experimental small-molecule agonist - danuglipron (PF-06882961) (19). Activation of GLP-1Rs by glucagon-like peptide-1 (GLP-1) stimulates insulin secretion and inhibits glucagon release, while delaying gastric emptying. The positive neuroendocrine effects of GLP-1R stimulation result in reduced food intake and improved satiety (19, 20). However, the main drawback of currently available injectable incretin therapies lies in patient adherence, their complex use, and side effects (19, 20, 21). Consequently, orally available small-molecule agonists with a good safety profile could offer greater therapeutic flexibility (19).

We first performed a deep learning-driven fragmentation of ruxolitinib and danuglipron, alongside probabilistic embedding to explore the chemical vicinity of their fragments. Subsequently, fragments were selected for new molecule *in silico* retrosynthesis using the breaking of retrosynthetically interesting chemical substructures (BRICS) algorithm (22).

This step was followed by virtual screening to explore binding energetics. The screening confirmed that most new candidate molecules had similar or improved binding compared to the parent molecule. Interestingly, new candidates demonstrated interactions that were not present in the original target molecule. In the case of danuglipron, our analysis hints at another key residue important for binding and molecule orientation that was not identified by previous studies (19). Top candidates were also highly unique when searched against known drugs. In parallel, new molecules were used to optimize a pharmacophore-based search, which helped identify two more molecules with markedly improved binding energetics. Thus, fragmentation could be part of the virtual screening pipeline, informing the search and early candidate selection as well as guiding a more focused optimization either through pharmacophore-based search or via candidate-driven library development.

We also aimed to bridge chemical and biological spaces by sampling the drug-target network induced by fragment mapping. Selecting fragment molecules and overlaying them with known biological targets of their parent compounds can be used to explore intricate interactomes. In particular, generated samples from ruxolitinib and danuglipron neighborhoods resulted in enriched clusters spanning multiple metabolic and immune nodes. These insights demonstrate that fragments in the latent space are organized by capturing features beyond the chemical space. Moreover, as our expressome analysis has shown, this strategy could be critical in establishing additional trajectories for drug repurposing and predicting side effects. For example, the ruxolitinib-induced network includes proteins responsible for sphingosine-related signaling which could be exploited in certain indications (23). This subcluster could potentially be suppressed by the downregulation of close neighbors, dependent on JAK2 interactions. Similarly, off-target effect prediction could offer better risk management for any drug discovery pipeline. The recent discontinuation of Pfizer’s danuglipron phase 3 clinical trial underscores the importance of such new methodologies in research and discovery (R&D) (24, 25). Notably, we also identified interactors that may have contributed to drug-induced liver injury observed in Pfizer’s trial.

Our findings established that fragmentation can aid in rational molecule design, using significantly smaller compound libraries compared to the current commercial practices. An in-depth look into the fragment-induced interactome and its integration with omics datasets can provide guidance for determining the therapeutic spectrum and off-target events. This is especially important for clinicians who must continuously adapt and expand treatment protocols to improve responsiveness and limit adverse effects.

## Methods

### Data and data preparation

Training data consisted of 11,548 small molecules that were in clinical development or had already been approved (26). This set was cleaned of duplicates and redundancies, resulting in 10,930 unique molecules. Testing and validation data (454 data points) were generated from the blood-brain barrier penetration (BBBP) dataset, after removing any overlaps with the training data (27). ZINC15 (>13 M compounds) (28) was employed for pharmacophore search. DrugBank (29) was used for similar compound search. Structures for the analysis were downloaded from the PDB database (30), namely GLP-1R (PDB ID: 7S15; cryogenic electron microscope resolved receptor complex with danuglipron analogue PF-06883365) and JAK2 (PDB ID: 6VGL; co-crystal structure with the type-I JAK inhibitor, ruxolitinib, bound to the JAK2 kinase domain in the active phosphorylated conformation). Drug-target search was performed with DGIdb (31) and the interactome parsing relied on the STRING database (v12.0) (32). KEGG database was used for additional pathway exploration (33). The expressome analysis included a bulk RNA sequencing study for ruxolitinib (GSE263709) (34) and an RNA sequencing of a preclinical mouse model fed a high-fat diet (HFD) to induce a metabolic syndrome for the treatment with semaglutide (GSE266899) (35).

### Bioinformatics, cheminformatics, and data analytics

Python (v 3.10) and R (v 4.4.1) were used as the programing environments. Python packages utilized included Numpy (v 2.2.4), pandas (v 2.2.3), seaborn (v 0.13.2), scikit-learn (v1.6), and matplotlib (v 3.10.1). R libraries employed for the analysis were STRINGdb (v2.16.4) and biomaRt (v2.60.1). RDKit (v 2.13) and Datamol (v0.12.5) facilitated compound analysis and preparation. Molecules were processed using Simplified Molecular Input Line Entry System (SMILES) encoding with implicit hydrogens. Compounds were fragmented using the BRICS algorithm (22) in combination with Bemis-Murcko scaffold identification rules (36). Cutoffs for fragment size were set between 3 and 18 atoms, inclusively. The BRICS algorithm (22) was used for in silico retrosynthesis of new compounds in the Datamol (v0.12.5) environment. Docking was performed with SwissDock Suite (37) and AutoDock Vina (v1.2.0) (38) with the following parameters for JAK2: box center: 28 - 10 – 52, box size: 22 - 22 – 22, sampling exhaustivity: 4, and GLP-1R: box center: 68 - 68 – 62, box size: 22 - 35 – 22, sampling exhaustivity: 4 . SwissSimilarity Suite was employed for a comprehensive compound search, comparing 2D fingerprints and electroshape between molecules of interest and DrugBank references (39). Compound absorption, distribution, metabolism, and excretion (ADME), as well as potential toxicity, were profiled with SwissADME Suite (40).

Pharmacophore search was performed with Pharmit against the ZINC15 database (41). PyMOL (Version 3.0 Schrödinger) and ChimeraX (v1.9; 42) analyses and visualization tools were used for interaction and surface visualization. Adaptive Poisson-Boltzmann Solver (APBS) was utilized for electrostatic potential mapping (43), while lipophilicity was determined based on molecular lipophilicity potential (MLP) principles (44). Both properties were visualized using the ChimeraX suite. Network analysis for fragments involved searching the STRING database (v12.0) with a minimum required interaction score = 0.4. Markov chain clustering (MLC) with inflation set to 3 was applied for interactor group identification. Enrichment parameters were set to false discovery rate (FDR)<=0.05, signal>=0.5, strength>=0.5, min. network count n=2 and background= whole genome (STRING database, v12.0). R2 Genomics Analysis and Visualization Platform (https://r2.amc.nl/) was used for expression profiling and analysis, following recommended parameters, setting FDR<=0.05 and absolute log2 fold change (log2FC)>=0.37.

### Deep learning

Positional processing and graph embedding was done via Molgraph (v 0.8.0). Deep learning framework was built with Tensorflow (v 2.15), Tensorflow probability (v0.22), and Keras (v2.13). Computational environment employed a Tesla T4 core with 16GB high-bandwidth memory (GDDR6). The network consisted of the encoder, the decoder, the prior set for regularization (cluster size: 3,5,10,15,20,25,30), and the integrated Gumbel softmax learning layer (Tau=0.25), which guided the appropriate selection of a GM layer. Latent space embedding dimensions were fixed at 32. Kullback-Leibler regularization was integrated with the probability layer with the following parameters: batch size=250, Monte Carlo sampling size=1000, test point reduction axis=0, weight=1/(batch size * sampling size)). Activation (where applicable) was ‘selu’ and the dropout rate was set to 0.2. Encoding and decoding loss was a custom root mean squared error (RMSE), which was also supplemented by Kullback-Leibler for detected clusters between the encoder and decoder weighted by 0.25. A detailed network schema is available in Supplementary Figure 2. The Adam optimizer with learning rate=1e-4 and weight_decay=True, and batch size set to 250 were used to train the model over 250 epochs with appropriate validation sets. Principal component analysis (PCA) was used for latent space visualization and dimensionality reduction after the data points had been normalized using StandardScaler (scikit-learn, v1.6).

## Results

### Graph-based probabilistic modeling revealed an intricate organization of the chemical space

A ChEMBL library comprising 10,930 compounds was filtered for training, with almost 6,000 compounds belonging to phase 2 clinical trials, and approved drugs included 2,336 therapeutics (Suppl. Fig. 1). For modeling purposes, we focused on the molecular features derived from SMILES (with implicit hydrogens) to characterize molecules using Molgraph (45). The number of atoms under these constraints ranged from 1 to 227. Similarly, molecules contained between 0 to 252 bonds. The average number of atoms per molecule was 26.23 (Suppl. Fig. 1). Training (n= 10,324) and validation (n=227) libraries were prepared by restricting fragment sizes between 3 and 18 atoms, inclusively. These readouts guided our fragmentation strategy to generate 23,314 molecules for training and 1,396 fragments for validation.

Instead of grouping entire compounds - which could limit the identification of therapeutically meaningful subgroups - the fragmentation approach enabled us to map the chemical space across different therapeutics (Fig. 1). Detailed mathematical reasoning can be found in Supplementary Document 1. By using Laplacian PE and GIN convolutional network integration, we were able to capture both local and global graph structures. The inputs are encoded with Laplacian PE, which combines molecule featurization with topological graph information. The model is trained to decode this input, as reconstructing graph structure directly has limitations in accuracy (46). GM embedding of molecular fragments provided a method to model complex spaces and to uncover probabilistic distributions of data points. Although determining cluster distributions might rely on different theoretical assumptions (i.e., Dirichlet processes, categorical selection, and others), we note that our latent space modeling supports more flexible cluster assignments with different underlying meanings. For our study purposes, each GM was governed by a predefined cluster number, selected from a discrete set that did not exceed the number of latent space dimensions and did not include the edge case of an isotropic Gaussian. The training converged with a 25-cluster GM model, achieving the overall loss after the 250^th^ epoch: 0.0153 and KL penalty: 0.00011819 (validation loss: 0.0193, KL validation penalty: 0.0160) (Suppl. Fig. 2).

Figure 2 illustrates how the fragments of our targets, namely ruxolitinib and danuglipron, are distributed in the learned latent space. It shows that simpler fragments cluster towards the top-right corner. Furthermore, the fragments evolve in complexity outwardly from this point. It is important to note that the latent space mapping is primarily governed by the researcher’s objectives; alternative approaches, such as kernel PCA, could be used to better visualize non-linear relationships. In specific cases — particularly for smaller size compounds — the entire molecular entity can be included in the mapping.

**Figure 2.**
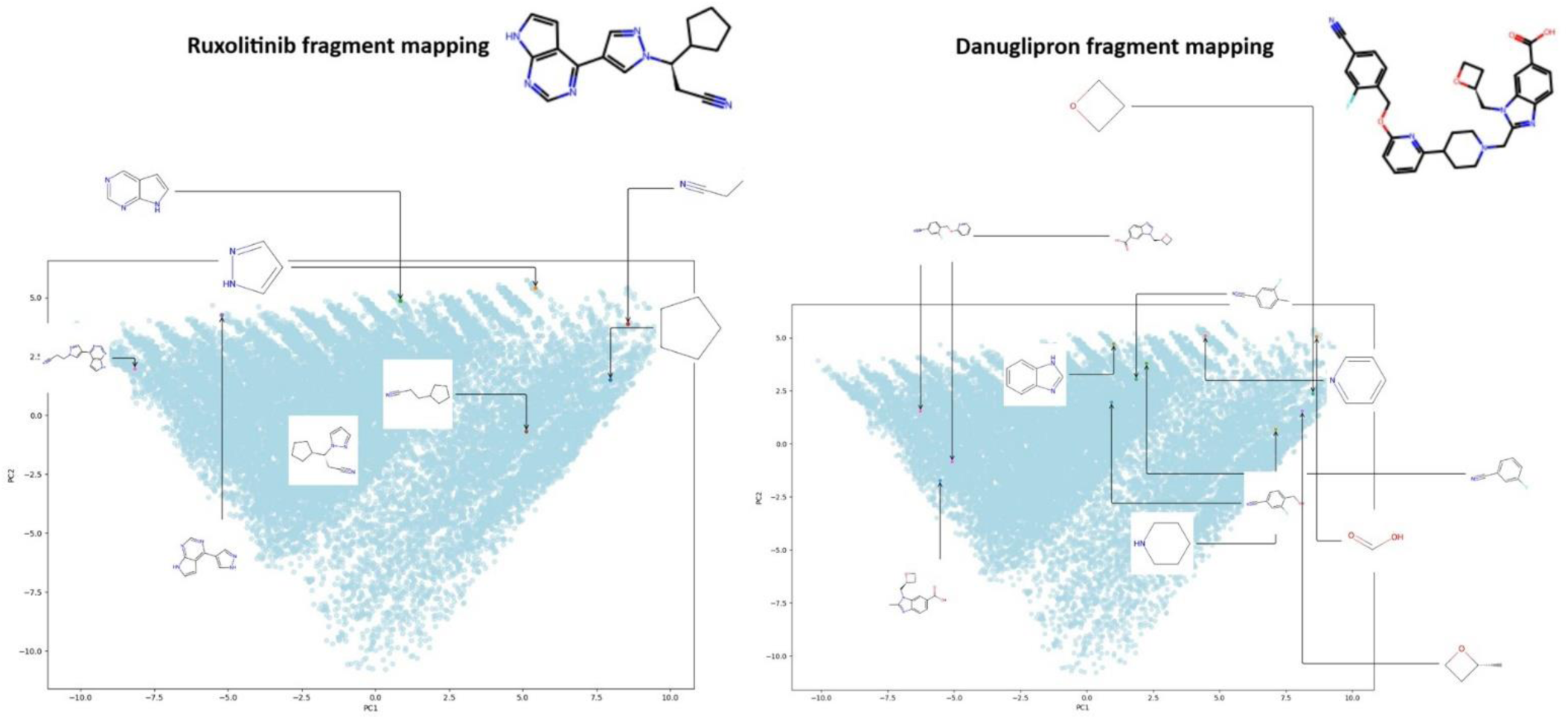
Ruxolitinib and danuglipron fragment mapping with the target compound displayed. PCA transformation is used for dimensionality reduction and better visualization of the latent space, showing the first two principal components.

### Fragment-guided molecule generation led to new and improved binding candidate detection

We turned our attention to the 10 nearest neighbors of every mapped fragment of ruxolitinib and danuglipron (assessed via mean squared error for distance). This analysis returned a diverse set of chemical entities for both drugs (Suppl. Fig. 3 and Suppl. Fig. 4). The neighboring fragments informed the selection of specific candidates for new molecule generation (Suppl. Document 2) that structurally and chemically shared at least several features with the parent molecule. Assembly was performed by iterating through all possible fragment binding arrangements under the BRICS algorithm (22). We then chose three candidates for JAK2 and GLP-1R based on the predefined criteria (Suppl. Document 2). Newly generated molecules were docked using AutoDock Vina (38) by setting the binding site, identified through crystallographic and cryogenic electron microscopy studies (JAK2 bound to ruxolitinib PDB ID: 6VGL; GLP-1R with danuglipron analogue PF-06883365 used for the binding site study PDB ID: 7S15). Thus, these experimental studies served as a reference point for the location, and the docking of ruxolitinib and danuglipron established a baseline for binding energetics (Table 1).

**Table 1.**
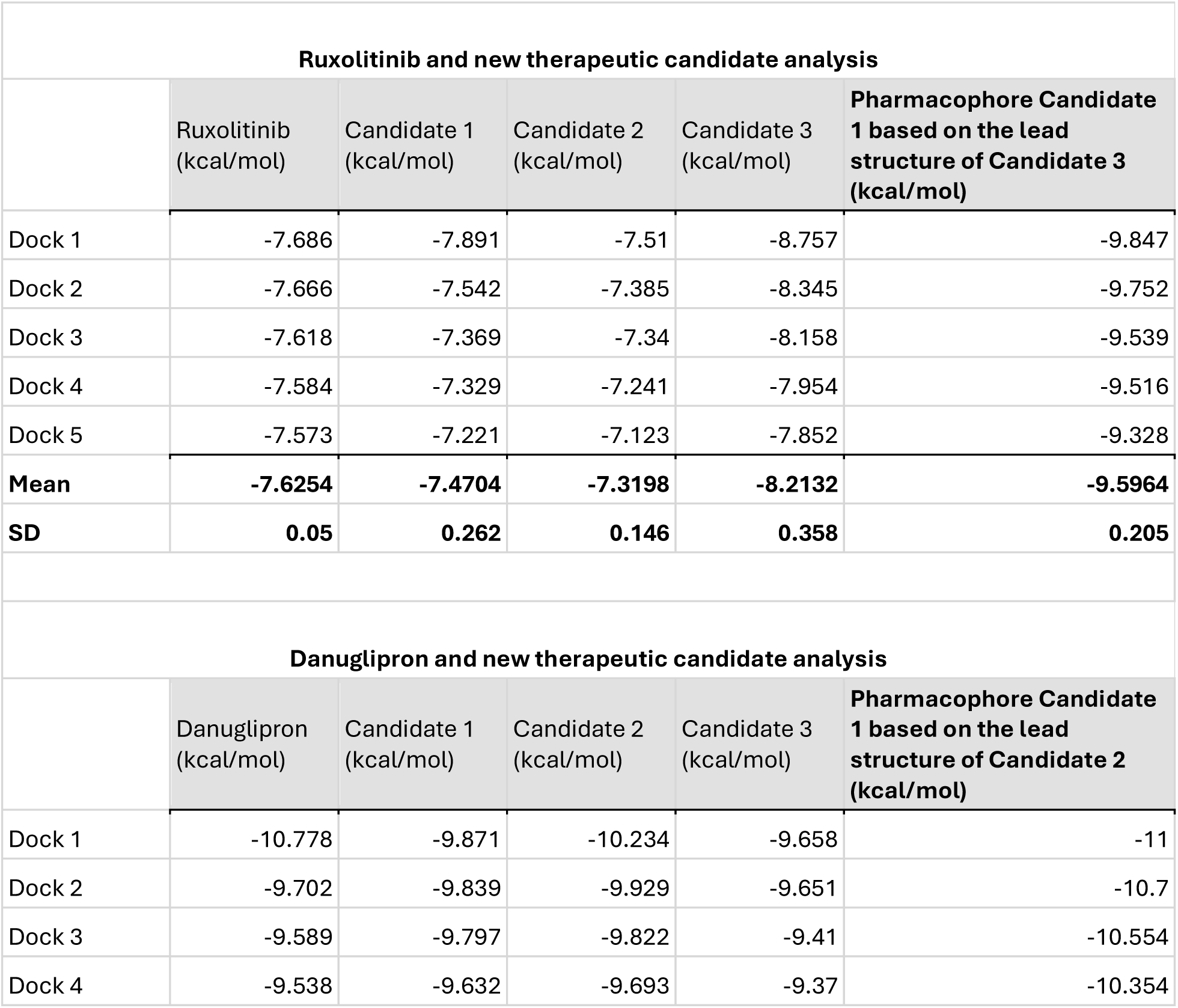
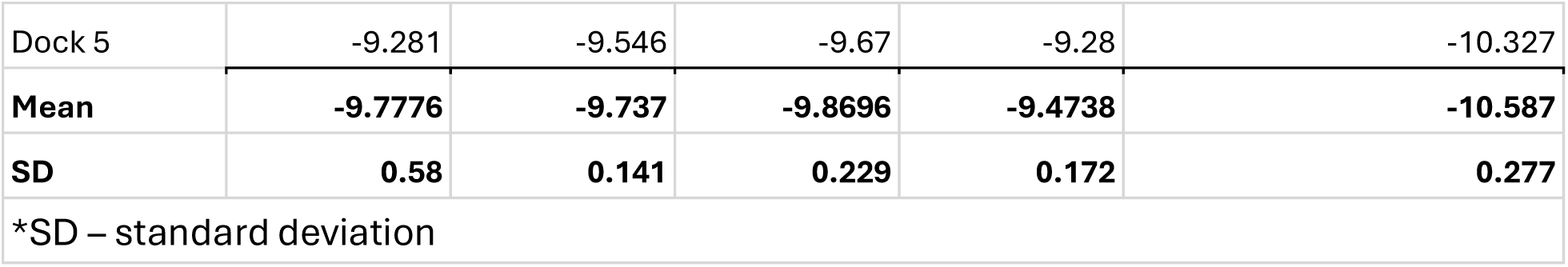
Docking results for compounds.

The docking summary (Table 1) reveals that the five best binding poses for ruxolitinib were close in energetics (SD: 0.05 kcal/mol), with the best binding pose scoring -7.686 kcal/mol. The crystallographically resolved structure of ruxolitinib was the closest to the fifth best pose (-7.573 kcal/mol). Candidates 1 and 2 exhibited more variation but had values comparable to the parent molecule. Interestingly, Candidate 3 demonstrated the best overall binding (top pose: -8.757 kcal/mol; mean: -8.2132 kcal/mol), while also showing the most variability (SD: 0.358 kcal/mol). In contrast, danuglipron varied the most in binding poses when compared to the other three candidates (SD: 0.58 kcal/mol), and it also had the best binding score of -10.778 kcal/mol. In addition, the resolved structure for GLP-1R with the danuglipron analogue mirrored the best binding pose model of the drug molecule. Other candidates selected for danuglipron optimization had lower variability, with Candidate 2 achieving an overall better binding score (mean: -9.8696 kcal/mol; SD: 0.229 kcal/mol).

The observed dynamics in binding energetics prompted us to investigate interactions of the top poses for the parent and candidate molecules (Table 1 and Fig. 3).

**Figure 3.**
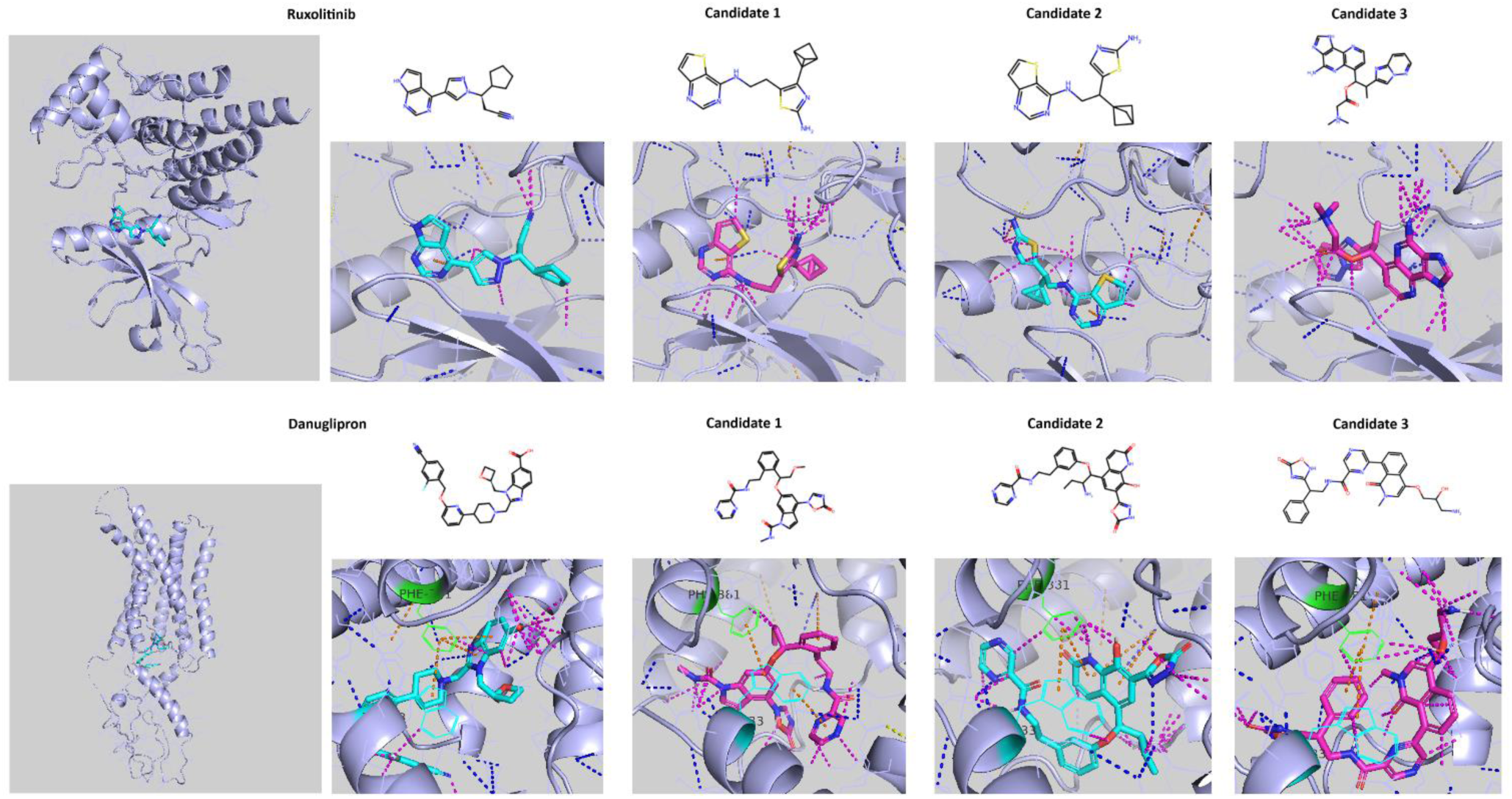
Docking results for the target and generated molecules, where the pose with the best docking score was analyzed. Protein structures include JAK2 (PDB ID: 6VGL) and GLP1-R (PDB ID: 7S15). Blue represents hydrogen bonds and polar bonds, yellow - ionic bonds, orange - all π interactions, magenta - all other interactions within 3.5 Å (PyMOL interaction graphics). Highlighted residues for danuglipron are PHE381 (green) and TRP33 (cyan).

As illustrated in Figure 3, ruxolitinib docks via several sidechain contact points and by forming a single π-coordination bond between the lysine residue (LYS882) and the pyrrolo[2,3-d]pyrimidine core. Other candidates have noticeably more anchoring points, including polar, non-covalent, and hydrogen bonds. Candidate 3 employs multiple contacts with side and main chains, which likely contribute to the observed binding strength. Danuglipron appears to have a complex coordination with phenylalanine (PHE381) and tryptophan (TRP33) via 1H-1,3-benzodiazole ring. The drug is further stabilized through polar and non-polar contacts with the side chains. For example, Candidates 2 and 3 share similar diverse interactions within the binding pocket; however, Candidate 1 no longer participates in π-coordination with PHE381(Fig. 3). Moreover, Candidate’s 1 pyrazine ring and TRP33 orient edge-to-face their π orbitals, and PHE381 maintains a strained coordination with TRP33 via parallel-displaced intrachain interactions.

Based on the overall binding performance, we selected Candidate 3 for ruxolitinib and Candidate 2 for danuglipron to search for similar drugs. We used a combination fingerprint (electrosphere and 3D fingerprint) to search DrugBank for any similar chemical entities.

However, none of the identified candidates exceeded a similarity score of 0.4 (Suppl. Fig. 6). Consequently, we further investigated whether we could exploit potentially promising interactions and build a pharmacophore for a large library search to identify better compounds.

We began by anchoring various pharmacophore features to the ruxolitinib Candidate 3 for the ZINC15 library search. This iterative process resulted in identifying an optimal pharmacophore structure that led to the hit compound ZINC5203833 (Fig. 4). The subsequent docking returned a markedly improved binding (best result: -9.847 kcal/mol, mean: -9.5964 kcal/mol, SD: 0.205 kcal/mol; Table 1). Analysis of APBS electrostatic and MLP surface revealed that Candidate 3 is seated deeply within the binding pocket, surrounded by the electron-rich surface with hydrophobic sites adjacent to methyl groups. The electrophilic and nucleophilic site distribution is mirrored by donor-acceptor sites on the pharmacophore. ZINC5203833 takes advantage of these contacts together with π-coordination (LYS882), resulting in a noticeably improved binding. We followed the same analysis steps for danuglipron Candidate 2. The molecule is positioned deep inside the binding pocket of the receptor that has both hydrophilic and hydrophobic sites (Fig. 4).

**Figure 4.**
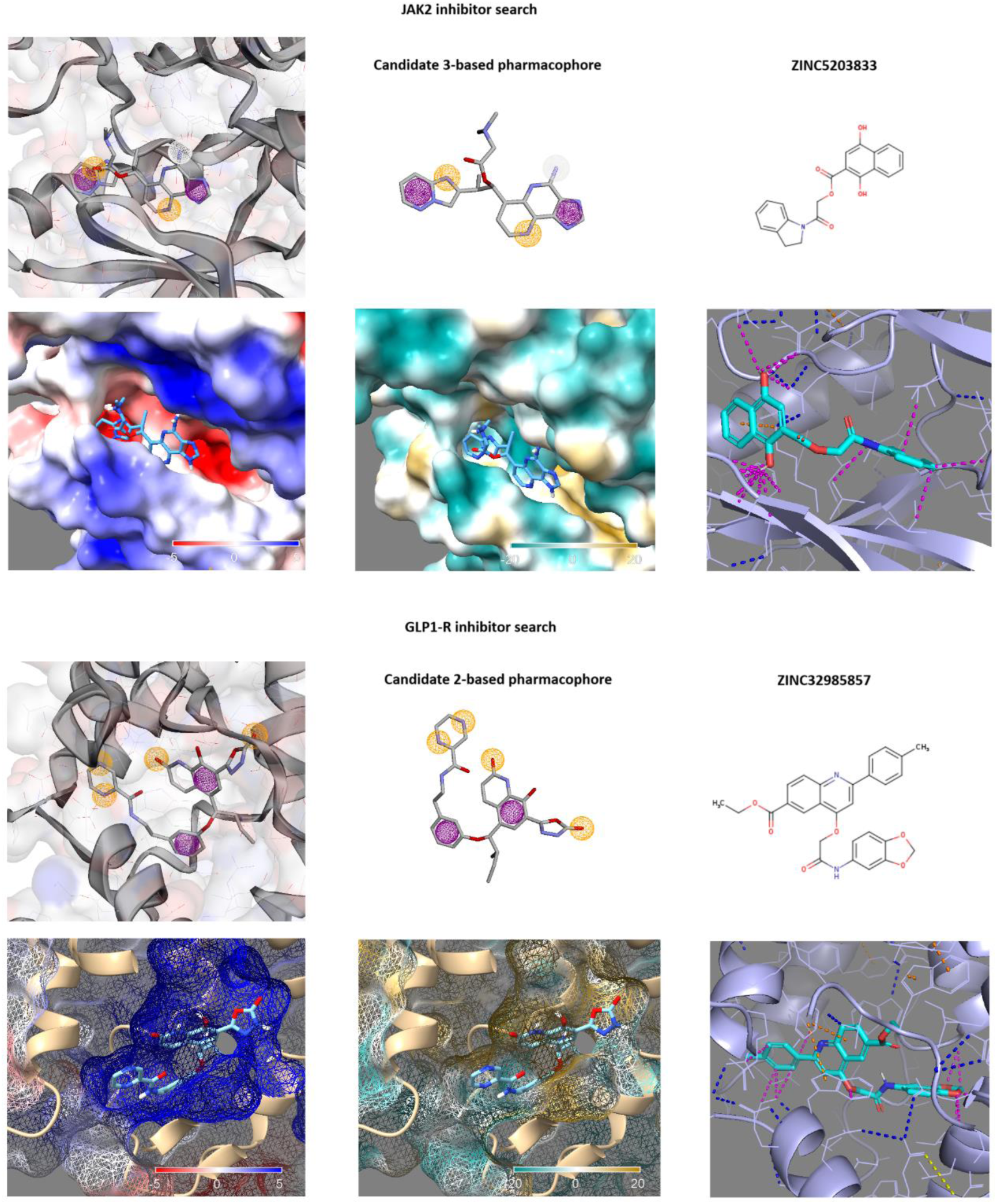
For each protein, the top three panels depict the pharmacophore model (with and without the binding site), where orange spheres (r=1 Å) depict hydrogen acceptors, white spheres (r=1 Å) are hydrogen donors, and magenta spheres (r=1 Å) represent aromatic groups (Pharmit visualization). The matched structure is displayed in the top-right corner for a given panel set. Bottom panels for each protein, from left to right, visualize Adaptive Poisson-Boltzmann Solver potential (APBS), with a range from -5 kT/e (red) to blue (+5 kT/e) and molecular lipophilicity potential (MLP), where dark cyan is the most hydrophilic site and dark goldenrod is the most hydrophobic area (ChimeraX visualization) for respective pharmacophore models. The last panel shows binding interactions for the identified molecule (blue represents hydrogen bonds and polar bonds, yellow - ionic bonds, orange - all π interactions, magenta - all other interactions within 3.5 Å) (PyMOL visualization). GLP-1R surface is shown as a mesh for better visibility of the buried compound.

Using the mapped pharmacophore for compound search returned ZINC32985857, which demonstrated the best binding energetics when compared to both the generated molecules and the parent compound (best result: -11.000 kcal/mol, mean: -10.587 kcal/mol, SD: 0.277 kcal/mol) (Fig. 4, Table 1). Notably, engagement with phenylalanine (PHE381) and tryptophan (TRP33) residues is also favored when orienting the compound in the constricted binding site (Fig. 4). This was also observed in danuglipron binding (Fig. 3). All compounds were analyzed for their ADME and toxicity profiles, which, combined with further lead studies and experimental testing, could aid in improved lead formulation (Supplementary Table 1).

### Fragment-guided interactome search identifies new interactors and networks

To gain insight into the interactome for ruxolitinib and danuglipron, we used a previously identified fragment neighborhood to map molecules to their parent compounds (PCA-based dimensionality reduction). Since fragments can have a varying number of associated parent molecules, it was important to normalize by randomly selecting only one compound per fragment. Alternatively, researchers can search for specific drugs that match fragments of interest for better lead optimization. These compounds created a search set for drug targets in the DGIdb database (31). For ruxolitinib, we identified 47 drug neighbors and their targets, above the set interaction threshold of 0.5 (Suppl. Table 2).

Using the STRING database, we mapped interactions for these proteins (minimum required interaction score=0.4), which returned a network of 45 nodes (PPI enrichment p-value: 6.77e-07) (Fig. 5). As can be seen, there is one major cluster (n=15, Cluster 1) with two prominent subclusters centered around KRAS (cancer metabolism and signaling) and JAK2 (immunological signaling). Noticeably, these proteins are involved in cross-cluster communication, either directly or through close network members, emphasizing overlaps for different biological processes (dashed lines, Fig. 5). For example, ADORA2A or adenosine A2a receptor is linked to Cluster 1 through FGFR1 (fibroblast growth factor receptor 1). The adenosine receptor is part of purinergic signaling pathways (Fig. 5, Cluster 2 enrichment). IL-7 receptor signaling is also in close proximity to JAK2, overlapping with multiple pathways and regulatory networks (Fig. 5, Cluster 1 enrichment). Clusters 5 and 7 provide further examples of metabolic networks that might span a spectrum of physiological functions (Fig. 5). Ghrelin (GHRL) is the ligand of growth hormone secretagogue receptor type 1 (GHSR), and they both play a role in satiety, intestinal transit, growth functions, as well as neuroactive signaling (KEGG database ID: hsa04080). Cluster 7 consists of sphingosine signaling associated members, S1PR1 and S1PR5, that play a role in multiple cellular processes. Despite the fact that the largest Cluster 1 dominates the network and the other groups consist of only 2 or 3 members, the significant enrichment suggests the possibility of a more complex interactome.

**Figure 5.**
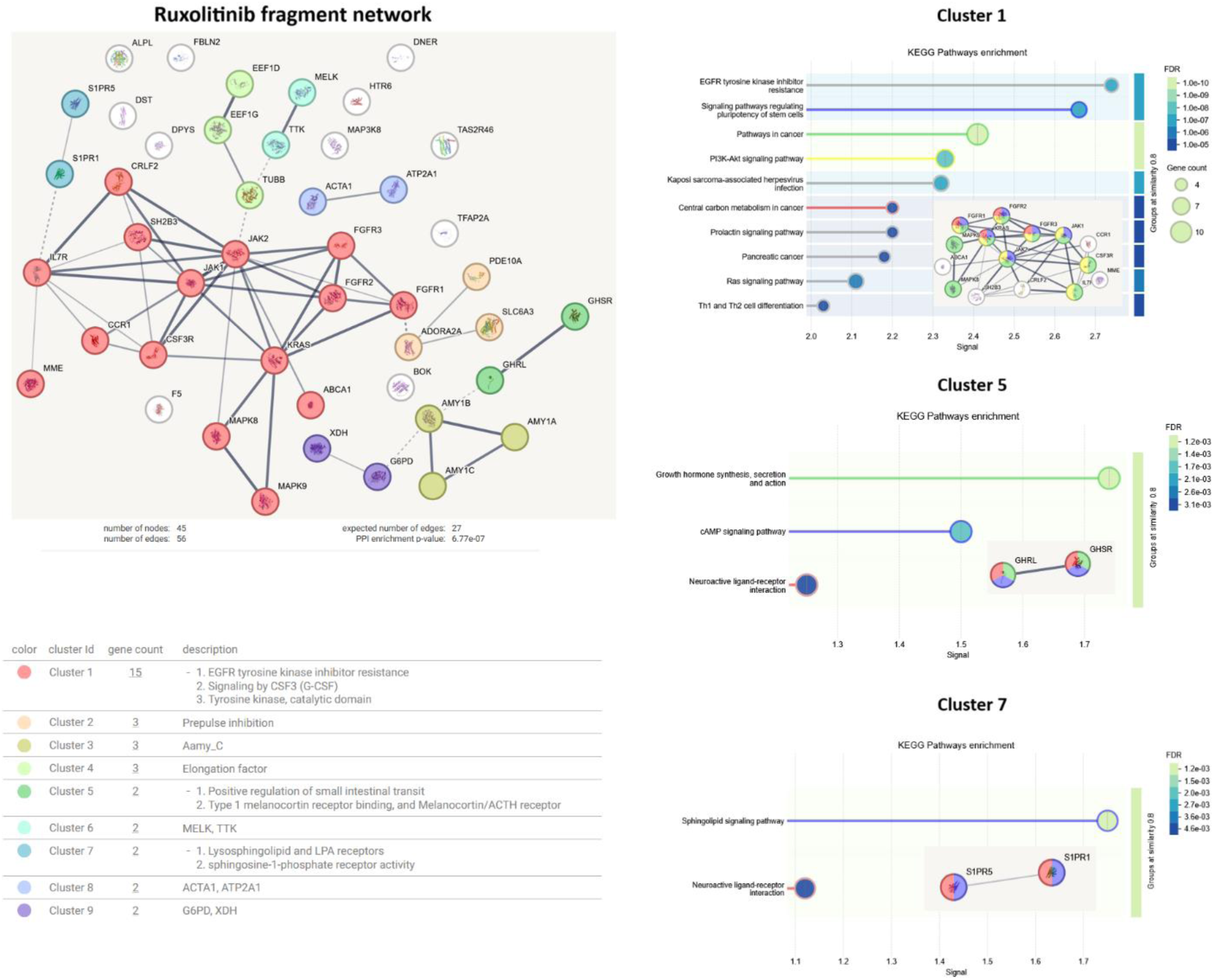
Ruxolitinib fragment neighborhood network and enrichment for selected biological processes and pathways. STRING database search: minimum required interaction score = 0.4, edges represent confidence. Ruxolitinib network of 45 nodes was clustered using the MCL algorithm with inflation parameter set to 3, where edges between clusters are shown with a dotted line. KEGG pathway enrichment is shown for individual clusters, along with the network fragment and highlighted pathways (enrichment: FDR<=0.05, signal>=0.5, strength>=0.5, min. network count n=2 background= whole genome; functional enrichment display: grouping by 0.8 similarity).

We also mapped interactions (minimum required interaction score=0.4) for the danuglipron neighborhood (n=62, DGIdb interaction threshold > 0.5; Suppl. Table 3), which resulted in a dense network of 57 nodes (PPI enrichment p-value: < 1.0e-16) (Fig. 6). The GLP-1R cluster (Fig. 6, Cluster 4) has several links with Cluster 1 that maps to a number of cellular functions and cancer metabolism pathways. The GLP-1R network is situated closely to 5-hydroxytryptamine receptor 2B (HTR2B), which is a serotonin receptor regulating a number of physiological functions. Similarly, G protein subunit beta 3 (GNB3) is another interactor of GLP-1R that has multiple metabolic functions. However, the largest cluster (n=10) of the studied protein set (Cluster 1) is associated with cancer and cancer metabolism pathways. For instance, the activation of the feline McDonough sarcoma-like tyrosine kinase (FLT3) pathway is essential for cell survival, cell proliferation, and differentiation of hematopoietic progenitor cells (47). The second-largest cluster (n=8) is part of the myosin network. This protein group is not only isolated from the rest of the clusters (no direct links at the set threshold) but it is also significantly enriched in cardiac health and homeostasis associated processes. Other smaller subgroups include chemokine receptors, solute carriers, ion channels, and proteasomal as well as cellular transport regulators, illustrating the broad connectivity of the danuglipron fragment network.

**Figure 6.**
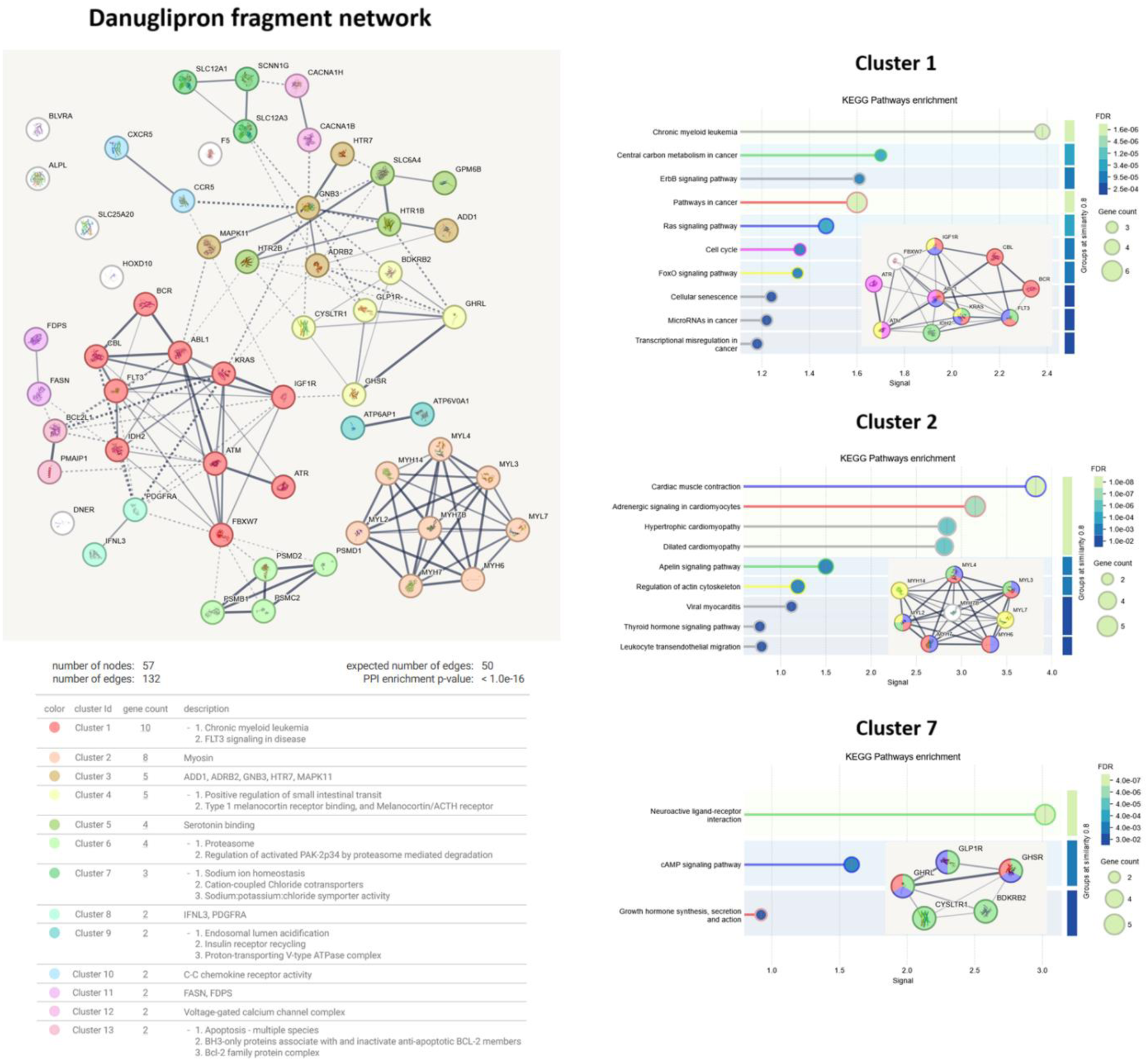
Danuglipron fragment neighborhood network and enrichment for selected biological processes and pathways. STRING database search: minimum required interaction score= 0.4, edges represent confidence. Danuglipron network of 57 nodes was clustered using the MCL algorithm with inflation parameter set to 3, where edges between clusters are shown with a dotted line. KEGG pathway enrichment is shown for individual clusters, along with the network fragment and highlighted pathways (enrichment: FDR<=0.05, signal>=0.5, strength>=0.5, min. network count n=2 background= whole genome; functional enrichment display: grouping by 0.8 similarity).

By capturing the interactome of ruxolitinib and danuglipron, we can establish signposts for potential modulation of other targets, which could be used for repurposing or adverse reaction prediction.

### Expressome analysis confirms a complex regulatory network for the selected targets

To complete our investigation of ruxolitinib and danuglipron networks, we performed an analysis of available drug screening studies.

We selected a ruxolitinib study on Philadelphia chromosome-like acute lymphoblastic leukemia (Ph-like ALL) to assess the identified network node responses to the treatment and disease perturbations. This study evaluated gene expression profiles in Ph-like ALL patient xenograft (PDX) models treated with control or ruxolitinib at different time points (GSE263709; 34). We performed a differentially expressed gene analysis using the R2 Platform between the control and the third, as well as the seventh treatment days for a drug-sensitive model (Fig. 7a). We found nine genes to be significantly changed from our network (Fig. 5), which spanned different regulatory nodes. For example, membrane metallo-endopeptidase (MME), maternal embryonic leucine zipper kinase (MELK), TTK protein kinase (TTK), tubulin beta class I (TUBB), and fibroblast growth factor receptor 1 (FGFR1) were upregulated after a three-day treatment. In addition, SH2B adaptor protein 3 (SH2B3), C-C motif chemokine receptor 1 (CCR1), ATP binding cassette subfamily A member 1 (ABCA1), and JAK2 were downregulated following the three-day treatment with ruxolitinib (Fig. 7a). When comparing these results with the differential expression in a seven-day model, the expressome profile changes: ABCA1 becomes upregulated, JAK2 returns to baseline, and a new gene, glucose-6-phosphate dehydrogenase (G6PD), becomes suppressed. Moreover, FGFR1, MME, SH2B3, and CCR1 retain their differential expression. Such fluctuations likely result from the initial network perturbation and subsequent adjustment within the disease model.

**Figure 7.**
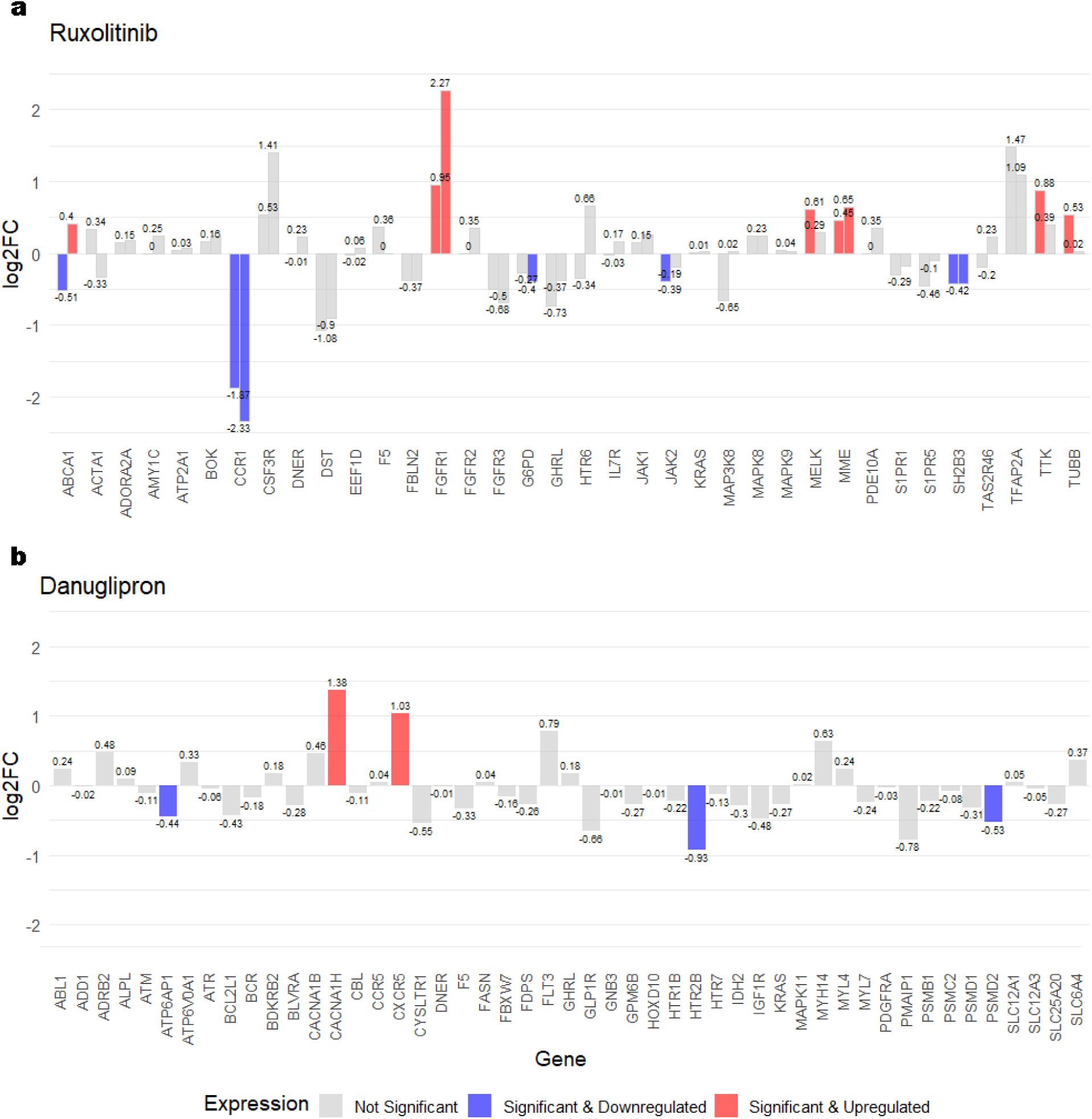
Differential expression (DE) profiles from external studies for assessing drug-induced perturbations within the fragment networks, reported as log2 fold change (log2FC). Only genes from networks with at least one non-zero log2FC are shown. (a) DE results comparing Day 3 (left bars) and Day 7 (right bars) after ruxolitinib treatment to control samples from Ph-like ALL PDX models. (b) DE results between semaglutide-treated (a GLP-1R agonist similar to danuglipron) and control samples from a high-fat diet mouse model.

In order to explore the danuglipron interactor network, we sought studies of similar profiles, as there were no publicly available datasets for this specific drug. We opted to investigate the gene expression perturbation in mice with induced metabolic syndrome that were treated with semaglutide - a GLP-1R agonist widely used in clinical practice (GSE266899; 35). Despite the study’s focus being pancreatic ductal adenocarcinoma, lesion sequencing provided additional insights into metabolic changes (Fig. 7b).

Specifically, proteasome 26S subunit ubiquitin receptor, non-ATPase 2 (PSMD2), ATPase H+ transporting accessory protein 1 (ATP6AP1), and HTR2B were significantly downregulated when exposed to the GLP-1R agonist. In contrast, C-X-C motif chemokine receptor 5 (CXCR5) and calcium voltage-gated channel subunit alpha-1H (CACNA1H) showed a marked increase in expression. While the sequencing study focuses on oncological pathology, semaglutide-induced alterations that can be mapped to our network underscore complex modulation across several regulatory hubs (Fig. 6 and 7b).

Even though these studies inherently had limitations and expression patterns are influenced by both pathology and tissue type, our findings reflect the need for more research on how drug-induced expression changes can affect therapeutic outcomes and influence side effect emergence.

## Discussion

In silico approaches in drug discovery have expanded the possibilities for new therapeutics identification. Nevertheless, there are a number of outstanding challenges for mapping the vast chemical space that by some estimates can even approach 10^100^ (48, 49). In particular, not only the amount of data, but also the ability to uncover meaningful interactions is crucial for discovery success. Thus, while most current methods put emphasis on activity, ADME, de novo design, and synthesis planning, relatively little attention has been given to fragment profiling (48). In the present work, we sought to redefine perspectives on fragment-guided drug characterization and new molecule generation. As case studies, we selected ruxolitinib and danuglipron based on their therapeutic targets and network modulation potential, which enables broader clinical impact. Our developed model aligns potentially meaningful chemical groups without the restriction of these fragments being ‘locked’ in a compound, which could bias the search. Moreover, our proposed strategy significantly reduced the size of the search space to only 10,324 molecules. This contrasts sharply with commercial and other discovery campaigns that rely on compound libraries ranging from millions to billions (6). By taking advantage of fragment-informed chemical space mapping, we were able to generate new compounds that showed improved binding when compared to the parent molecule. In addition, fragment sampling for danuglipron yielded novel molecules that revealed a previously unrecognized residue as important for binding and coordination. To complete the in silico pipeline, we incorporated biological space mapping to uncover new interactors that could be used for other modulator discovery or off-target effect prediction. Consequently, our analysis not only provided further insights into targeted therapy development but also outlined gene sets of particular importance for pathophysiology management.

To achieve our study goals, we introduced a new deep learning model in which the network’s architecture is dynamically learned to identify the optimal latent space encoding through a GM layer (Fig. 1, Suppl. Fig. 2). Leveraging the expressivity of graph encoding (50), the first layer of molecular mapping was realized through Laplacian PE and GIN convolution layers (Suppl. Fig. 2). Specifically, the PE and graph convolutions captured both chemical and topological interactions, which allowed for richer fragment feature integration. Since our focus was on therapeutic entities, we demonstrated that this approach, even with a relatively small training set (∼10,000 molecules), can lead to a discovery of completely new and structurally diverse compounds with little similarity to any marketed drugs (Suppl. Fig. 6). This highlights the importance of developing new methods for drug design that do not solely rely on large chemical libraries (48, 51).

We tested newly designed compounds by benchmarking them against parent molecules, namely ruxolitinib and danuglipron. Ligand docking is still challenging, owing to the fact that there are many considerations, ranging from flexibility to the scoring function’s accuracy (52). However, we wanted to explore how conformational ‘snapshots’ could accommodate molecules and whether this information could offer new insights. Thus, we employed two resolved structures for JAK2 and GLP-1R in which the parent molecule was bound, aiding with the docking site’s identification. Our newly identified molecules for both target proteins demonstrated a number of new interactions that were not present in the original drug (Fig. 3). Consequently, these interactor groups could be used to establish modifications necessary for improved binding and new therapeutics design (Fig. 3, Table 1).

To further characterize novel compounds, we started our analysis with ruxolitinib (Suppl. Fig. 4, Suppl. Document 2). Candidate 3 has several key anchoring points via dimethylamine. The same candidate also employed a number of polar and non-polar contacts via the three-ring heterocycle (Fig. 3). From an electrostatic and lipophilicity perspective (Fig. 4), it is evident why these contacts are important in accommodating a molecule in the JAK2 binding pocket. Subsequently, these observations facilitated a pharmacophore search, which resulted in an even stronger in silico binding profile (ZINC5203833) (Fig. 4). It is worth noting that new compound generation in combination with pharmacophore-based search can narrow down likely leads and guide targeted library search. In the case of danuglipron (Suppl. Fig. 5, Suppl. Document 2), the binding pocket is much more constrained with limited degrees of freedom for molecule binding. Our analysis confirmed earlier findings that tryptophan (TRP33) is required for danuglipron signaling (19) as binding is coordinated through this residue. However, we also identified phenylalanine (PHE381) which mediates anchoring for danuglipron in combination with TRP33, which was not highlighted previously (19; Fig. 3). After comparing docking results of the candidates, we selected Candidate 2 for pharmacophore search, as it exhibited not only a completely different docking pose, but also a more intricate quadruple coordination system involving TRP33, PHE381, LYS197, and its quinoline derivative core (Fig. 3). This approach returned the ZINC32985857 molecule (Fig. 4, Table 1) with a markedly improved binding and the same coordination system of PHE381, TRP33, and compound’s quinoline-like core. Evidently, both GLP-1R residues are necessary for optimal molecule accommodation within a binding pocket (Fig. 4). Thus, we not only identified new candidates with improved binding when benchmarked against the parent molecule but also showed that an additional residue within the GLP-1R binding pocket is essential for binding. Importantly, such an exploratory refinement of candidate-to-pharmacophore demonstrates that we can efficiently identify novel therapeutic candidates and probe molecules to refine potential leads.

For the next step in our analysis, we investigated whether we could extend the chemical fragment space to the interactome to gain deeper insights into drug interactions, repurposing, or on/off-target effects. To achieve this, we analyzed the close neighborhood of our target fragments and sampled associated drugs. These drugs were utilized to capture additional targets for network construction. The mapped targets showed a significant enrichment for various interactome clusters (Fig. 5 and Fig. 6). This suggests that fragments carry their own ‘functional meaning’ that extends to biological processes. Additionally, functional similarities between fragments could be leveraged for therapeutic optimization, drug repurposing, or predicting potential side effects. Similarly, new functionalities or biomarkers could be explored by identifying novel interactors. Thus, ruxolitinib and danuglipron networks allowed us to better understand the therapeutic profiles of these drugs and to theorize about newly designed modulator effects.

JAK2 inhibition involves a varied set of signaling events. Exploring beyond the nearest neighbors, such as KRAS, we discovered several key metabolic trajectories (Fig. 5). For example, ADORA2A stands out as a close neighbor to the JAK2 cluster, where it interacts with FGFR1 (Fig. 5, Cluster 2 enrichment). Moreover, FGFR1 links ADORA2A and the adenosine signaling with KRAS and the JAK2 subclusters via the Rap1 pathway (KEGG database ID: hsa04015), which translates into cellular adhesion, migration, polarity, and survival. Targeting adenosine has been examined as a treatment strategy to block adenosine production and its receptor binding in oncological disorders (53, 54). However, recent clinical studies on JAK2 inhibition for anti-tumor effects have not established a link between JAK2/STAT3 and adenosine modulation effects, despite its potential effects in modifying tumor microenvironment and inducing immune reactions (55). Our expressome analysis revealed that FGFR1 was significantly upregulated when ALL xenografts were treated with ruxolitinib (Fig. 7). This could alter regulatory dynamics such that, in oncological indications, FGFR1 might contribute to resistance together with sustained stimulation of ADORA2A. Nevertheless, in neurological disorders the FGFR1-ADORA2A axis could promote neuroplasticity and offer alternative treatment avenues (56, 57). To our knowledge, there have been no reports in the clinical drug development setting identifying this link or exploring the potential of multi-modulation and different therapeutic outcomes, depending on the disease. Another member of the JAK2 cluster, SH2B3, regulates IL-7 receptor signaling (58) and links inflammation with hypertension (59). Notably, JAK2 has been implicated in salt-sensitive blood pressure and experimental studies established that JAK2 inhibition with ruxolitinib decreases pulmonary arterial pressure, restoring cardiac index (60, 61). Additionally, ruxolitinib treatment in the ALL model downregulated both JAK2 and SH2B3, which points to another therapeutically relevant signaling branch (Fig. 7a). IL-7R connects this interactor group with an adjacent small cluster consisting of sphingosine-1-phosphate receptors that play a role in regulating inflammatory processes (62). Furthermore, experimental studies have indicated that S1PR1 activates the JAK2/STAT3 pathway via a complex feedback loop (63). Thus, these observations underscore a broader impact of this network on several physiological functions and a complex interplay between different regulators, which could help establish new therapeutic strategies. Correspondingly, interactome analyses can flag potential side effects or parameters requiring clinical monitoring. For example, an initial drop in ABCA1 expression followed by upregulation (Fig. 7a) could have an effect on high-density lipoprotein (HDL) levels. A study in mice demonstrated that ruxolitinib decreases cholesterol efflux, accelerating atherosclerosis (64, 65). Therefore, more research is needed to define clinically relevant alterations that could influence treatment protocols or pose a higher risk if other comorbidities are present. Taken together, a fragment-driven chemical and biological space mapping can help us better understand the drug’s role in both therapeutic and adverse outcomes.

Turning our attention to the GLP-1R network, we observe similarly enriched sub-clusters. The GLP-1R cluster (Cluster 4) has several links with Cluster 1 that maps to a number of cellular functions and cancer metabolism pathways. This is particularly noteworthy since the effects of GLP-1R agonists on cancer biology have recently gained more attention (66). Our fragment-based network mapping highlights these links through several network trajectories, such as serotonergic, chemokine, and GPCR signaling (Fig. 6). For example, serotonin signaling has been implicated not only in immune cell regulation but also in oncological and metabolic disorders (67). Semaglutide-induced HTR2B downregulation in the pancreatic ductal adenocarcinoma model (Fig. 7b) (35) mirrors other studies demonstrating that HTR2B inhibition can suppress colorectal cancer tumor growth (68).

The complex effects of serotonin are evident through other studies of HTR2B antagonism, which attenuated high-fat diet-induced visceral adipose tissue inflammation, hepatic steatosis, and insulin resistance (69). Another therapeutically relevant group is formed by myosin (Cluster 2), which includes the apelin signaling pathway (Fig. 6). Given that apelin is involved in the pathogenesis and protective effects in many diseases, such as coronary heart disease, heart failure, diabetes, and cancer, this network arrangement hints at the modulation potential of a number of parallel pathways (KEGG database ID: hsa04371) (70). Furthermore, GLP-1R has been documented for its cardioprotective effects; however, the exact mechanisms, for example, in heart failure with preserved ejection fraction (HFpEF), remain to be elucidated (71). A good example for such a therapeutic scope expansion is also the N-type calcium channel (CACNA1B) (Cluster 12) which contributes to both cardio-and neuroprotective effects through the cervical ganglion (SCG) of the autonomic nervous system (72). The role of SCG in cardiovascular diseases has been documented (73, 74), and further studies are necessary to establish CACNA1B modulatory effects on cardiac tissue. One member of the voltage-gated calcium channels, CACNA1H, was shown to be markedly upregulated in the pancreatic cancer study. This could provide further evidence for potential anti-cancer effects of GLP-1R agonists (35) (Fig. 7b) as CACNA1H inhibitors were found to reduce apoptosis induced by the ER stress, and in oncological scenarios it could increase treatment sensitivity (75). Downregulation of PSMD2 and ATP6AP1 has a similar anti-proliferative effect (Fig. 7) (76, 77). As can be seen, by juxtaposing other medications, we can map a broader spectrum of beneficial functions across several therapeutic categories for GLP-1R. Currently, adverse effects, such as nausea, vomiting, or other gastrointestinal issues, associated with GLP-1R agonists have not been traced to specific pathway overstimulation, outside of the proposed GLP-1R+ neuron activation (25, 78). In particular, none of the preclinical and clinical danuglipron studies have incorporated genetic profiling to address the complex signaling of GLP-1R (19, 25, 79, 80). This lack of a more in-depth drug action characterization is evident across other GLP-1R clinical studies (25). In our analysis, GLP-1R is in close proximity to amiloride-sensitive sodium channel subunit gamma (SCNN1G), which has been implicated in several disorders (81). In particular, SCNN1G mediates luminal sodium and water diffusion through the apical membrane of epithelial cells, thereby regulating electrolyte and blood pressure homeostasis (81). Interestingly, it also has a role in taste perception (82). As a result, SCNN1G and other solute carriers in the network should be examined for adverse effects relating to hypertension, kidney tissue damage, gastrointestinal irritation, and electrolyte changes (81). Moreover, our network analysis points to several pathways that can perturb liver metabolism leading to tissue injury (Fig. 6). For example, the FoxO signaling pathway (KEGG database ID: hsa04068) is known to regulate liver stress response and hepatic homeostasis (83). Several up- and down-stream members of this network, namely ATM serine/threonine kinase or Ataxia-telangiectasia mutated (ATM), insulin-like growth factor 1 receptor (IGF1R), and KRAS, connect with the GLP-1R cluster (Fig. 6). Other examples include C-C chemokine receptor type 5 (CCR5), which has been reported to play a role in chronic liver injury development (84). We also reported an upregulation of CXCR5 which could signal B cell infiltration, and, in the liver, this can lead to a pro-inflammatory state (Fig. 7b) (85). These observations - in light of Pfizer’s failed phase 3 clinical trial of danuglipron due to drug-induced liver injury - further emphasize missed opportunities in the drug’s development (24). Therefore, integrating genetic parameters into clinical endpoint assessment could prevent clinical trial failure by tailoring studies for populations most likely to benefit from the treatment while also improving predictions for adverse effects (25).

Both the JAK2 and GLP-1R networks share six common members (i.e., Delta and Notch-like epidermal growth factor-related receptor (DNER), GHRL, GHSR, alkaline phosphatase (ALPL), factor V (F5), KRAS), accentuating their connection to pro-inflammatory states and metabolic regulation. Notably, the pharmacological intervention of the JAK-STAT pathway has emerged as a promising therapeutic approach for the treatment of metainflammation. This condition results in chronic and low-grade systemic inflammation and is characteristic of obesity, type 2 diabetes, and other disorders (86). Therefore, multi-modulatory approaches for complex diseases are increasingly being considered to achieve optimal treatment outcomes (86). For example, an experimental diabetes study in mice demonstrated that liraglutide (GLP-1 analog) inhibited the JAK2/STAT3 pathway (87). Other noteworthy regulatory networks in this context involve the neurological axis connecting both JAK2 and GLP-1R. Within the ruxolitinib network, SLC6A3, which is a dopamine transporter (Fig. 5, Cluster 2), links the JAK2 subnetwork with dopaminergic signaling. Dopamine is indispensable in regulating T cells, B cells, natural killer (NK) cells, macrophages, and dendritic cells (DCs) (88, 89). Consequently, dopamine agonists have been explored as a therapeutic strategy for rheumatoid arthritis (90). Similarly, the GLP-1R cluster extends into serotonergic signaling (Fig. 6, Cluster 5), which also modulates many immune and metabolic responses (91, 92). Given the discussed examples, it becomes evident that managing complex diseases could benefit from expanding the scope of therapeutic targets and their neighborhood mapping. Moreover, the growing number of multimorbidities will require improving personalized care to balance drug action (93). As a result, our introduced integrative approaches may offer better therapy matching and modeling of side effects.

## Conclusion and future directions

The growing burden of complex diseases underscores the need for improved therapeutic solutions that employ diverse and targeted modulation strategies. While the advent of deep learning has significantly advanced drug discovery, current strategies are often limited to sifting through large volumes of chemical data without optimizing alternative methods for exploration. Consequently, enhancing drug discovery methods and characterization would significantly improve how we develop and use therapeutics in clinical settings. Our work not only offers a new perspective on targeted therapy development but also introduces an interactome mapping strategy, which takes advantage of molecular fragment profiling. We also developed a new self-adjusting neural network for this purpose that helped discover novel therapeutic candidates with improved binding.

Furthermore, fragment-guided chemical space exploration enabled us to explore new metabolic trajectories. Our discussed clinical examples showcase how these insights could further guide drug repurposing efforts and advance the prediction of side effects. This is particularly important information for clinicians, who continuously refine treatment protocols for better outcomes and the minimization of adverse effects. Importantly, future research should expand on deep learning modeling methodologies as well as explore implications of broader network perturbations. Therefore, our work maps the first critical steps for this endeavor and highlights promising new avenues for drug discovery and clinical intervention development.

### Limitations

Fragment-based analyses currently have no comparable models or established benchmarks, which future research should address. While our study focused on the chemical space mapping and bridging it with the interactome using in silico and structural benchmarks, the next experimental step would be to perform targeted screens of new molecules to explore structure-activity relationships. Finally, it is important to stress that crystallographic or other structural studies explore proteins and ligands in conditions that may not replicate interactions occurring in vivo (i.e., minor or other binding poses could also exist). Thus, additional in vitro and in vivo characterization studies in combination with molecular modeling should follow to ensure robust lead candidate optimization.

## Supporting information

Abbreviations

Figure and table descriptions

## Data and Software Availability

Data are available from public resources as referenced in the Methods section. Algorithmic and mathematical implementations are further described in Supplementary materials. Code can be made available upon a reasonable request.

## Supporting Information

Mathematical outline for the model (Supplementary Document 1, PDF); Additional information for investigated compounds (Supplementary Document 2, PDF); Compound features (Supplementary Figure 1, PNG), Figure visualizing the model (Supplementary Figure 2, PNG), Training process outputs (Supplementary Figure 3, PNG), Relevant fragments for ruxolitinib (Supplementary Figure 4, PNG), Relevant fragments for danuglipron (Supplementary Figure 5, PNG), Similar compound search results (Supplementary Figure 6, PNG); Compound chemical profile summary (Supplementary Table 1, Comma Separated Values Source File); Ruxolitinib interactome selection (Supplementary Table 2, Comma Separated Values Source File); Danuglipron interactome selection (Supplementary Table 3, Comma Separated Values Source File)

## Acknowledgement section

This research received no external funding.

