## Supplementary material for "Fragment-Guided New Therapeutic Molecule Discovery and Mapping of Clinically Relevant Interactomes": Abbreviations

Å - angstrom; unit of length, equal to $10^{-10}$ meter

ADME - absorption, distribution, metabolism and excretion

APBS - adaptive Poisson-Boltzmann solver potential

DE – differential expression of genes

KL - Kullback–Leibler divergence

kT/e - the thermal voltage; a measure of the average kinetic energy of charge carriers in a material due to thermal motion

log2FC - log2 fold change; a measure for differential gene expression

MCL - Markov clustering

MLP - molecular lipophilicity potential

PCA – principal component analysis

PDX - patient-derived xenograft
