## Supplementary material for "Fragment-Guided New Therapeutic Molecule Discovery and Mapping of Clinically Relevant Interactomes": Figure and table descriptions

**Supplementary Figure 1**. Clinical phase and feature distribution plots for a 10,930 compound library. Phase assignment is based on a ChEMBL classification, where Phase -1 signifies an unknown status for a drug (i.e., its status cannot be accurately determined), Phase 0.5 represents compounds in early Phase 1 clinical trials. Phases 1-3 include compounds under clinical investigation, and Phase 4 is an approved marketed drug. Molecular feature histograms depict distributions for the number of aromatic rings, cycles, heteroatoms, bonds, and atoms. For modelling purposes, only the explicit atoms from SMILES encoding were counted.

**Supplementary Figure 2**. Network schema outlining the encoder and decoder structure.

**Supplementary Figure 3**. Model performance for training and validation sets.

**Supplementary Figure 4**. Ruxolitinib fragments and top ten nearest molecules based on the mean squared error as a distance measure.

**Supplementary Figure 5**. Danuglipron fragments and top ten nearest molecules based on the mean squared error as a distance measure.

**Supplementary Figure 6**. Similar compound search using a combination score of 3D fingerprint and electrosphere to search DrugBank.

**Supplementary Table 1.** ADME profiling for studied compounds (SwissADME suite).

**Supplementary Table 2.** Ruxilitinib and ruxolitinib fragment-based network where drug-target interactions were mapped from DGIdb. Interaction score was filtered based on the set threshold (>0.5), including the compound of interest regardless of the sore.

**Supplementary Table 3.** Danuglipron and danuglipron fragment-based network where drug-target interactions were mapped from DGIdb. Interaction score was filtered based on the set threshold (>0.5), including the compound of interest regardless of the sore.
