## Supplementary document 1 for "Fragment-Guided New Therapeutic Molecule Discovery and Mapping of Clinically Relevant Interactomes"

### Mathematical outline for the model

#### 1 Introduction

Each fragment is studied from a graph-theory perspective, where we define a graph as G = (V, E) with V being the set of nodes (or atoms) and E - the set of edges (or bond types). Given a graph G=(V, E) with $n$ vertices, its adjacency matrix A is an $n \times n$ matrix where $A_{ij} = 1$ if a bond exists between atoms $i$ and $j$; otherwise $A_{ij} = 0$. Laplacian PE is used to update fragment features before passing them into GIN convolutional and GM layers.

The objective of the variational autoencoder is to reconstruct the latent space encoding to match the input information. This is achieved using Bayes' law. The input data and the latent variable are denoted by $x$ and $z$, respectively. To model the latent space, we also define a categorical variable, $y$, representing the data point's membership of a Gaussian cluster. There are $K$ GM models, such that $k \epsilon \{1, \ldots, K\}$ indexes the GM model, which is used for latent space encoding. In the present work, both the approximate posterior $Q_{\phi_k}(y_k, z_k|x)$ and the latent space distribution defined by the prior $P(z_k)$ are Gaussian mixtures, where $k$ indexes a specific GM model from the set and denotes its associated parameters. Note that the multinomial distribution:

$$y_k \sim Q_{\phi_k}(y_k|x) \tag{1}$$

represents the cluster assignment probability for every data point in a latent space for a specific GM model, where $\phi_k$ represents parameters learnt by the model. $Q_{\phi_k}(z_k|y_k, x)$ represents a distinct Gaussian:

$$Q_{\phi_k}(z_k|y_k, x) = \mathcal{N}(\mu_{z_k}(x, y_k), \sigma^2_{z_k}(x, y_k)) \text{ (2) and categorical } k \sim P(k|x) \tag{3}$$

Our work expands on previous models (Varolgüneş et al., 2020) by allowing the model architecture to dynamically adjust and select the optimal number of clusters (or distribution $y_k$). Gumbel Softmax trick is used to guide the distribution selection process using a differentiable function. The probabilities of cluster assignment are learned through $P(k|x)$, which serves as a probability of selecting a specific Gaussian mixture model. Specifically,

$$P(k|x) = \frac{exp(log(\pi_k)+g_k)/\tau}{\Sigma_{j=1}^{k} exp(log(\pi_j)+g_j)/\tau} \tag{4}$$

so that $\pi_k$ – the probability of a specific Gaussian mixture model (logits) are encoded by learning from the inputs and $g \sim Gumbel(0, 1)$. $\tau$ sets the temperature allowing to regulate entropy of the outputs. Combining these steps results in the inference model

$$Q_{\phi_k}(y_k, z_k|x) = Q_{\phi_k}(z_k|y_k, x)Q_{\phi_k}(y_k|x) \quad (5)$$

The generative part of the model operates on the following probabilities:

$$P(y_k) = \text{Uniform}(\frac{1}{n_k}) \quad (6)$$

where $n_k$ is the number of clusters for a learnt selection of a GM model (eq. 3)

$$P_{\theta_k}(z_k|y_k) = \mathcal{N}(\mu_{z_k}(y_k), \sigma^2_{z_k}(y_k)) \quad (7)$$

and

$$P_{\theta_k}(x|z_k) = \mathcal{N}(\mu_x(z_k), \sigma^2_x(z_k)) \quad (8)$$

or the joint probability of reconstruction

$$P_{\theta_k}(x, z_k, y_k) = P_{\theta_k}(x|z_k)P_{\theta_k}(z_k|y_k)P(y_k) \quad (9)$$

(assuming conditional independence for $P_{\theta_k}(x|y_k, z_k) = P_{\theta_k}(x|z_k)$, where $\theta_k$ denotes parameters learnt by the decoder (10)). The resulting loss function can be summarized $\theta_k$ defines parameters leant by the decoder:

$$\mathcal{L}(\phi, \theta, k; x) = \mathbb{E}_{Q_{\phi_k}(y_k, z_k|x)}[log\frac{P(y_k)}{Q_{\phi_k}(y_k|x)} + log\frac{P_{\theta_k}(z_k|y_k)}{Q_{\phi_k}(z_k|y_k, x)} + logP_{\theta_k}(x|z_k)] \quad (11)$$

where the first part of the expectation represents the entropy, the second part is the regularization, and finally the reconstruction-based loss. For model learning purposes, we write an equivalent loss function for the predicted outputs $\hat{x}$:

$$\mathcal{L}(\phi, \theta, k; x) = RMSE(x, \hat{x}) + wKL_{reg}(d\_clusters||e\_clusters) \quad (12)$$

The root mean squared loss (RMSE) controls for appropriate fragment encoding and decoding. Weighted ($w$=0.25) $KL_{reg}$ helps to maintain the stability of the embedding space to appropriately learn the cluster number (e_clusters – encoded cluster distribution, d_clusters – decoded cluster distribution).
