## Supplementary document 2 for "Fragment-Guided New Therapeutic Molecule Discovery and Mapping of Clinically Relevant Interactomes"

**Target: Ruxolitinib** (JAK1/JAK2 inhibitor ), 6VGL

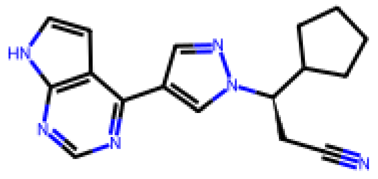

Ruxolitinib

**Generated candidates (fragment latent space search, BRICS algorithm)**

1. SMILES: Nc1nc(C2C3CC2C3)c(CCNc2ncnc3ccsc23)s1

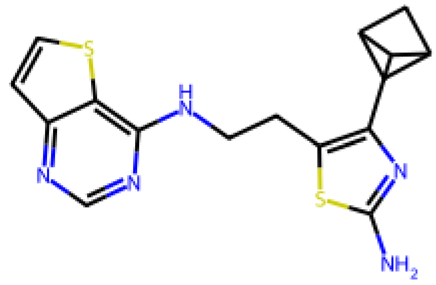

2.SMILES: Nc1ncc(C(CNc2ncnc3ccsc23)C23CC(C2)C3)s1

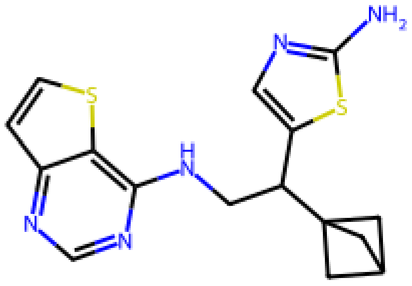

3. SMILES: CC(c1cn2ncccc2n1)C(OC(=O)C[NH+](C)C)c1ccnc2c1nc(N)c1nc[nH]c12

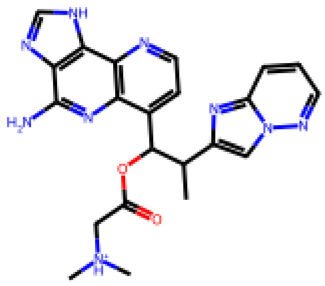

**Pharmacophore candidates based on Smiles 3**

**1. Pharmacophore candidate, ZINC5203833**

**SMILES:**O=C(OCC(=O)N1CCc2ccccc21)c1cc(O)c2ccccc2c1O

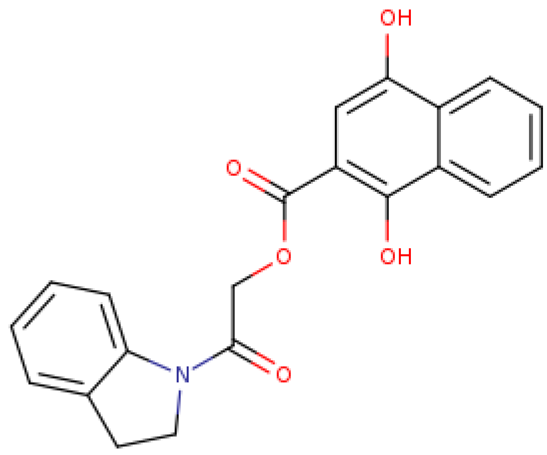

**Target: Danuglipron** GLP1 agonist small molecule (PDB entry 7S15)

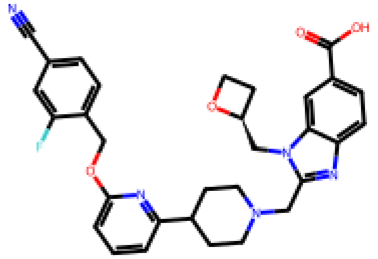

Danuglipron

**Generated candidates (fragment latent space search, BRICS algorithm)**

1. SMILES: CNC(=O)n1ccc2c(-n3cnc(=O)o3)cc(OC(COC)c3ccccc3CCNC(=O)c3cnccn3)cc21

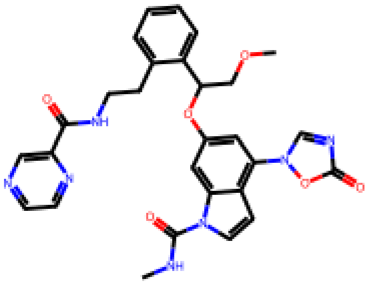

2. SMILES: CCC(N)C(Oc1cccc(CCNC(=O)c2cnccn2)c1)c1cc(-c2n[nH]c(=O)o2)c(O)c2[nH]c(=O)ccc12

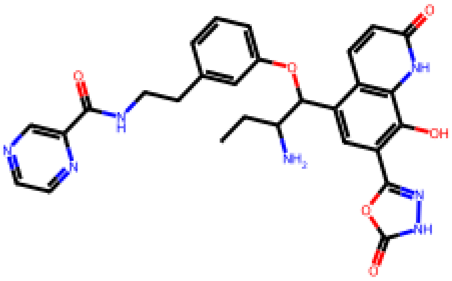

3. SMILES: Cn1cc(OCC(O)CN)c2cccc(-c3cncc(C(=O)NCC(c4ccccc4)c4nc(=O)o[nH]4)n3)c2c1=O

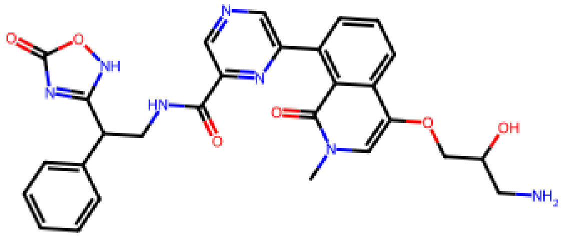

Top lead based on smiles 2 pharmacophore **[ZINC32985857]**

SMILES: CCOC(=O)c1ccc2nc(-c3ccc(C)cc3)cc(OCC(=O)Nc3ccc4c(c3)OCO4)c2c1

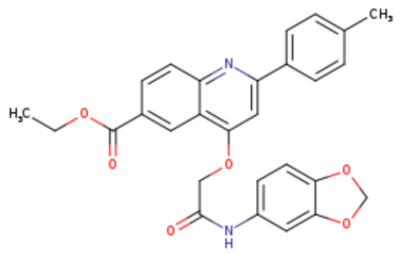
